# Topological Entanglement in Intrinsically Disordered Proteins: Sequence, Structural, and Functional Determinants

**DOI:** 10.64898/2026.03.23.713674

**Authors:** Wangfei Yang, Henry Silvernail, Debasis Saha, Eleni Panagiotou, Wenwei Zheng

## Abstract

Intrinsically disordered proteins (IDPs) populate heterogeneous conformational ensembles that are difficult to characterize using conventional structural descriptors. As a result, it remains unclear which ensemble features meaningfully connect sequence composition to biological function. Here, we employ entanglement-based measures derived from knot theory to provide complementary insight into IDP organization. Using the human IDRome database, we analyze two continuous entanglement descriptors, the writhe and the second Vassiliev invariant (*V*_2_), across more than 28,000 simulated disordered sequences. We show that these entanglement measures exhibit structured, low-dimensional variation across the database and display distinct relationships with sequence composition and ensemble geometry. Writhe primarily reflects compaction-dependent coiling tendencies that are largely recoverable from coarse sequence and structural features, whereas *V*_2_ captures higher-order topological organization that is less predictable from simple metrics. Embedding the resulting distribution features reveals functionally enriched regions of entanglement space, and ortholog simulations demonstrate that these signatures are evolutionarily conserved. Together, these results establish entanglement as a biologically relevant dimension of IDP organization and provide a rigorous, complementary framework for linking its sequence, ensemble structure, and molecular function.

## Introduction

Intrinsically disordered proteins (IDPs) constitute a broad class of biomolecules that perform essential regulatory and signaling functions without adopting a single well-defined tertiary structure.^1^ Unlike folded proteins, whose structures are encoded by a unique energy minimum,^2^ IDPs populate heterogeneous ensembles of conformations whose properties are determined by sequence-encoded biases in charge, hydropathy, and aromatic patterning.^3–5^ This structural plasticity enables IDPs to mediate diverse biological processes,^6^ from molecular recognition to the formation of biomolecular condensates.^7,8^

A central challenge in understanding IDPs lies in deciphering the relationship between sequence composition, the resulting conformational ensemble, and biological function. One possible approach is to infer function directly from sequence-derived properties such as amino acid composition, charge patterning, or hydropathy profiles. Recent large-scale analyses of IDP sequence grammars have demonstrated that purely sequence-based approaches can successfully recover functional regularities within disordered regions.^9–11^ However, this strategy remains fundamentally constrained by the enormous conformational freedom and the dynamic, heterogeneous nature of disordered ensembles. Even minor sequence or context perturbations can shift the populations of many interconverting substates across a wide energy landscape,^12,13^ producing subtle yet functionally important changes that are difficult to resolve without explicit structural or ensemble-level information. As a result, an intermediate, ensemble-based representation of IDPs is often necessary to bridge the gap between sequence and function.

Although numerous computational and integrative approaches can now generate IDP ensembles with reasonable accuracy,^14–24^ the ability to extract meaningful structural insights ultimately depends on how these ensembles are interpreted. A broad set of structural features has therefore been developed to describe disordered chains at multiple levels of resolution. Global descriptors such as the radius of gyration, end-to-end distance, and asphericity capture overall chain compaction and shape, and have been especially useful for linking sequence patterning to liquid-liquid phase separation propensities.^25–27^ Pairwise measures, including contact probabilities, inter-residue distances or distance distributions, characterize amino acid specific interactions that underlie context-dependent functions.^13^ Local metrics, such as secondary-structure propensities and solvent accessibility, reveal sequence-encoded biases in short-range organization, which often preconfigure regions for motif-specific interactions.^28^

Despite the breadth of available descriptors, it remains fundamentally unclear which structural features are actually relevant for biological function. Different molecular activities likely depend on distinct aspects of IDP organization, some driven by global chain compaction, others by specific long-range contacts, and still others by subtle local biases. Yet evolution does not tune sequences with respect to radius of gyration, contact probabilities, or secondary-structure propensities; these metrics reflect what we are able to measure or compute, not necessarily the variables that matter the most. Consequently, functional behavior may emerge from higher-order properties of the ensemble that are not captured by conventional local, pairwise, or global features. This motivates the need for complementary representations, such as topological or entanglement-based measures to quantify local and global topological complexity,^29^ that integrate information across scales and describe how the chain organizes in three-dimensional space. Unlike standard geometric descriptors, topological measures quantify the degree to which the chain winds, threads, or entangles, a global property that can influence segment accessibility, rearrangement dynamics, and even interaction pathways. These descriptors provide a natural way to capture global correlations that emerge from collective fluctuations, offering structural information that is inaccessible to traditional metrics. If such topological features are biologically relevant, they should be reflected in functional specialization and potentially conserved through evolution.

IDPs and their conformational ensembles can be modeled mathematically by open curves in three-dimensional space that do not intersect. The distinct topologies of simple closed curves in space are classified in knot theory. Via closure approximation schemes, knot theory has been productively applied in the folded-protein context, where persistent knots and related topological motifs have been used to analyze folding pathways and stability.^30–32^ However, these approaches provide classification and not quantification metrics. The only metric until recently that applied to quantify structural complexity of proteins was the Gauss linking integral in the form of writhe and linking number. ^33–35^ More recently, topological descriptors have begun to appear in studies of disordered systems. Local writhe analyses applied to short fragments have revealed transient coiling in highly dynamic IDPs,^36^ and LASSO entanglement metrics have been used to characterize threading events when forming transient contacts in disordered regions.^37^ In addition, Lang and colleagues recently introduced a “link-node” approach, systematically computing the Gauss linking number between sub-chains to identify persistent physical links in IDPs, and showed that entangled regions correlate with reduced local motions.^38^ A novel mathematical framework introduced in^39,40^ has further enabled the construction of new metrics for quantifying global topological structural complexity in proteins. Topological measures have since been applied to aggregated or fibrillar states of disordered proteins, where knot analyses help describe the complex entanglement patterns that emerge during assembly.^41,42^

However, it remains unclear whether topological organization in IDP ensembles carries biological meaningful information, particularly in function and evolution. While previous studies have quantified writhe-like measures in disordered chains,^33,36^ they have largely focused on geometric characterization and have not systematically examined whether these topological descriptors align with biological functions or evolutionary conservation. Moreover, higher-order topological metrics such as the second Vassiliev measure (*V*_2_)^40,41,43^ have not been explored in the context of IDPs. This gap highlights the need for a broad, quantitative analysis that connects topological organization to biological function and evolutionary constraints.

In this work, we investigate whether topological measures in IDP ensembles reflects biologically relevant features of function and evolution. Using conformational ensembles generated from molecular simulations of diverse IDP sequences in the IDRome database, ^44^ we quantify chain entanglement through two continuous topological observables: the writhe and *V*_2_. We first characterize how these quantities describe the structural organization of disordered ensembles and how they relate to sequence-derived properties and conventional structural descriptors. We then assess whether variations in topological organization are associated with particular molecular functions and whether these signatures are conserved across orthologous sequences. We find that both writhe and *V*_2_ exhibit well-defined, low-dimensional organization across the IDRome dataset, display distinct sequence and structural determinants, and importantly, align with functional classifications and evolutionary signatures that are not captured by conventional descriptors. Together, these analyses establish topological organization as an informative and previously underexplored bridge linking sequence, conformational ensemble, and biological function in disordered proteins.

### Interpreting structural entanglements of intrinsically disordered proteins

To investigate the topological organization of disordered proteins on a large scale, we utilized the IDRome database, which contains more than twenty thousand human intrinsically disordered sequences and their associated molecular trajectories.^44^ For multiple conformations within the conformational ensemble of each sequence, we computed two continuous topological descriptors, writhe and the second Vassiliev invariant (*V*_2_), which quantify distinct aspects of spatial self-entanglement (see Methods). As shown on the left side of Fig. 1A, writhe measures how much a chain coils around itself and with what handedness. Intuitively, it counts the signed self-crossings of the curve in a projection, so segments that wrap around each other in a right-handed sense contribute positively, while left-handed wrapping contributes negatively. Writhe is sensitive to the overall chirality of pairwise strand crossings. In contrast, *V*_2_ detects higher-order entanglement patterns arising from the global structure of a protein and is less sensitive to local geometrical changes. Its sign does not capture chirality, but reflects distinct characteristics of global topologies. As a result, configurations with similar overall coiling or even identical writhe can nevertheless differ in their global topological organization, which *V*_2_ resolves.

**Figure 1:**
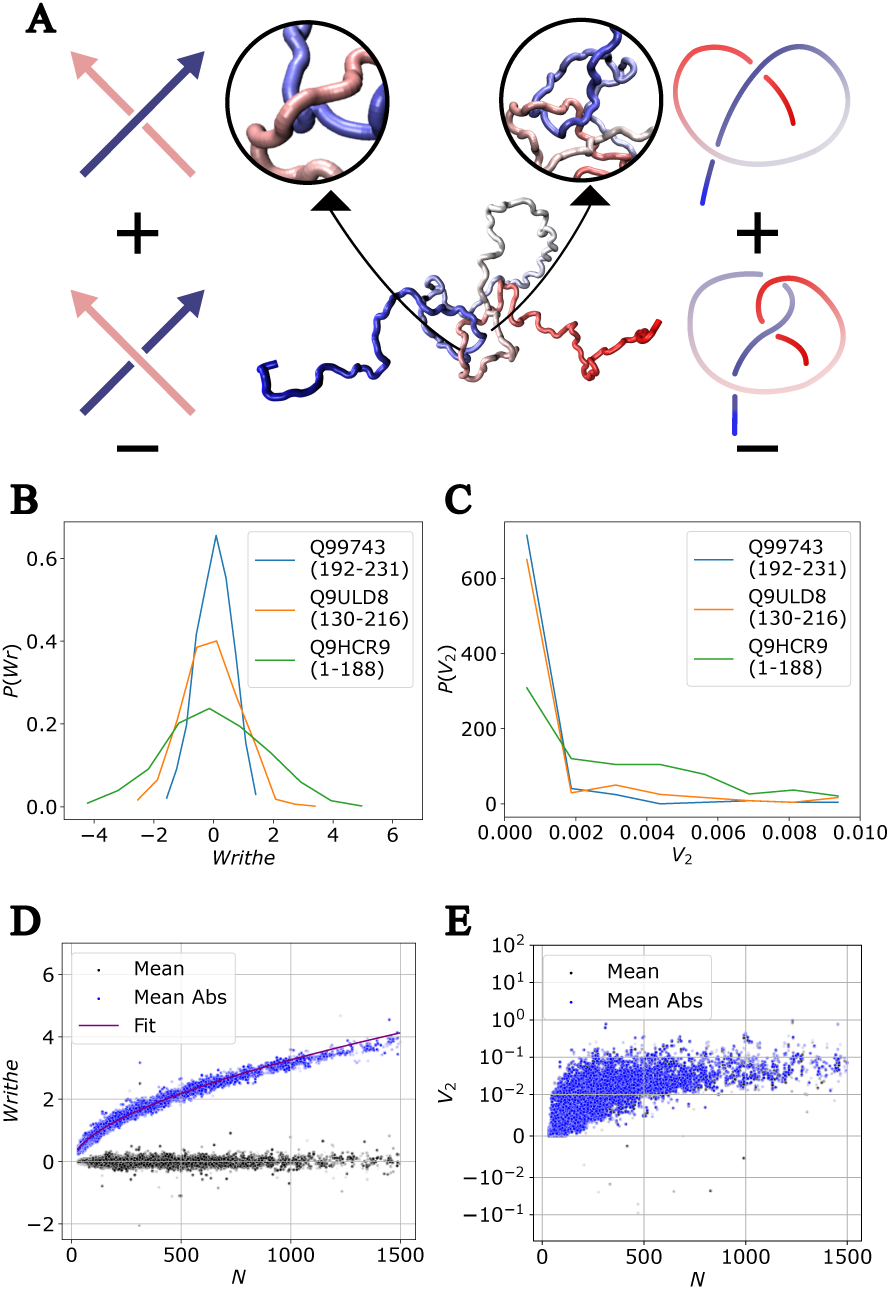
Topological characterization of conformational ensembles of intrinsically disordered proteins in the IDRome database.^44^. (A) Schematic illustration of the calculation of writhe and *V*_2_ from polymer conformations. (B) Representative probability distributions of writhe and (C) *V*_2_ for selected sequences. (D) Mean and mean absolute writhe as a function of chain length (*N*). The purple line shows a power-law fit of the form 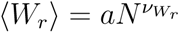. (E) Mean and mean absolute *V*_2_ as a function of chain length (*N*).

The writhe was computed for all 1000 conformations per sequence available in IDRome. In contrast, the calculation of *V*_2_ is substantially more computationally demanding. We therefore performed a systematic convergence analysis with respect to the number of conformations included in the averaging (see Methods for technical details). As shown in Fig. S1, *V*_2_ converges reliably when using 200 conformations per sequence. Based on this analysis, we adopted these parameters, which provide stable and sequence-independent estimates of *V*_2_.

Using these procedures, we obtained distributions of writhe and *V*_2_ values for each of the 28,058 sequences in the IDRome database. These distributions reflect the intrinsic variability of topological features within each ensemble. As shown in Fig. 1B, writhe exhibits a broad and nearly symmetric distribution centered near zero, indicating that IDPs frequently adopt conformations that coil in both right-handed and left-handed senses. In contrast, *V*_2_ displays an asymmetric distribution that is skewed toward positive values (Fig. 1C). This positive bias suggests that transient self-contacts in IDPs tend to form specific global topological structures. Configurations that generate negative *V*_2_ typically require specific arrangements (Fig. 1A), which are statistically less likely even in a random coil. A similar positively skewed *V*_2_ distribution has also been reported in previous studies of Tau filaments^41^ and random walks.^45^ Importantly, the width and shape of these distributions vary substantially across sequences, reflecting differences in both topological propensity and conformational heterogeneity. This variability highlights a fundamental distinction between disordered and folded proteins. Whereas folded proteins are characterized by a single dominant structure, IDPs sample a broad spectrum of self-entangled geometries. Accordingly, the full distributions of writhe and *V*_2_, rather than their ensemble averages alone, contain additional information that we exploit in the analyses that follow.

To examine how topological descriptors depend on chain length, we analyzed the scaling behavior of writhe across the full IDRome dataset (Fig. 1D). The mean absolute writhe increases systematically with sequence length, while the signed mean writhe shows no measurable chain length dependence. This difference reflects the underlying symmetry of the writhe distribution (Fig. 1B). Because writhe values span both positive and negative orientations, the mean absolute value grows simply because longer chains have more opportunities to coil in either direction, whereas the signed mean remains near zero. The mean absolute writhe therefore captures the intrinsic magnitude of self-entanglement, independent of cancellation between positive and negative contributions.

Having established the mean absolute writhe as the relevant chain-length dependent quantity, we next quantified its scaling behavior. Following standard approaches in polymer physics used for observables such as the radius of gyration,^46,47^ we fit the chain-length dependence of the mean absolute writhe to a power law of the form 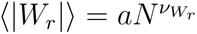, where *a* is a global prefactor and 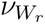 is the scaling exponent, as shown in Fig. 1D. Using this shared reference scaling relation, we convert the absolute writhe of each conformation into an effective writhe scaling exponent (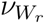). Conformations with larger exponents 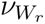 correspond to more pronounced or more frequent entanglement, whereas smaller exponents indicate conformations that are less entangled than expected for their chain length. Conceptually, 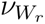 plays a role analogous to the polymer scaling exponent *ν* extracted from the radius of gyration (*R_g_* ∼ *N^ν^*), but it characterizes how topological self-entanglement, rather than spatial size, accumulates with chain length. We return to the relationship between these two scaling exponents in a subsequent section. The distribution of this writhe scaling exponent 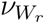, rather than the raw writhe, is carried forward in all subsequent analyses. In contrast to writhe, neither the ensemble-mean nor the mean absolute values of *V*_2_ shows a strong dependence on chain length across IDRome (Fig. 1E), we retain the raw *V*_2_ distributions without applying a chain-length normalization.

### Sequence correlates of entanglement metrics

In folded proteins, strong tertiary packing and specific side-chain interactions typically restrict the conformational ensemble to a single native topology, so topology is largely determined by the folded state. IDPs, however, do not adopt a single stable fold; instead their structural properties emerge statistically from an ensemble of conformations. In this ensemble, therefore, sequence-dependent biases, such as charge patterning or hydrophobic clustering, can systematically shift the probability of particular self-entanglement motifs. Accordingly, we ask to what extent do 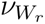 and *V*_2_ reflect primary-sequence features in IDPs? Establishing such sequence–topology links is a prerequisite for evaluating whether observed topological tendencies are biologically meaningful or amenable to design.

To evaluate how primary sequence influences topological organization, we computed pairwise correlations between sequence descriptors and 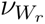 - or *V*_2_-based ensemble statistics across the IDRome database (Fig. 2A,B). Spearman’s rank correlation coefficient was used to capture monotonic relationships, including nonlinear dependencies, between sequence features and topological metrics. On the sequence side, we included chain length *N*, compositional fractions (charged, aromatic, and hydrophobic residues) and patterning measures such as *κ*,^3^ sequence charge decoration (*SCD*)^4^ and sequence hydropathy decoration (*SHD*).^5^ A full list and definitions of these sequence features are provided in Table S1. For each topological descriptor, we examined twelve distribution-level characteristics, including the mean, mean absolute, median (*Med*), standard deviation (*σ*), skewness (*γ*_1_), kurtosis (*γ*_2_), inter-quartile range (*IQR*), mean absolute deviation (*Mad*), and several extremal statistics, which together capture different aspects of the shape and breadth of the writhe and *V*_2_ distributions (Table S2). This framework enables a systematic assessment of which aspects of sequence architecture align with distinct statistical signatures of topological entanglement.

**Figure 2:**
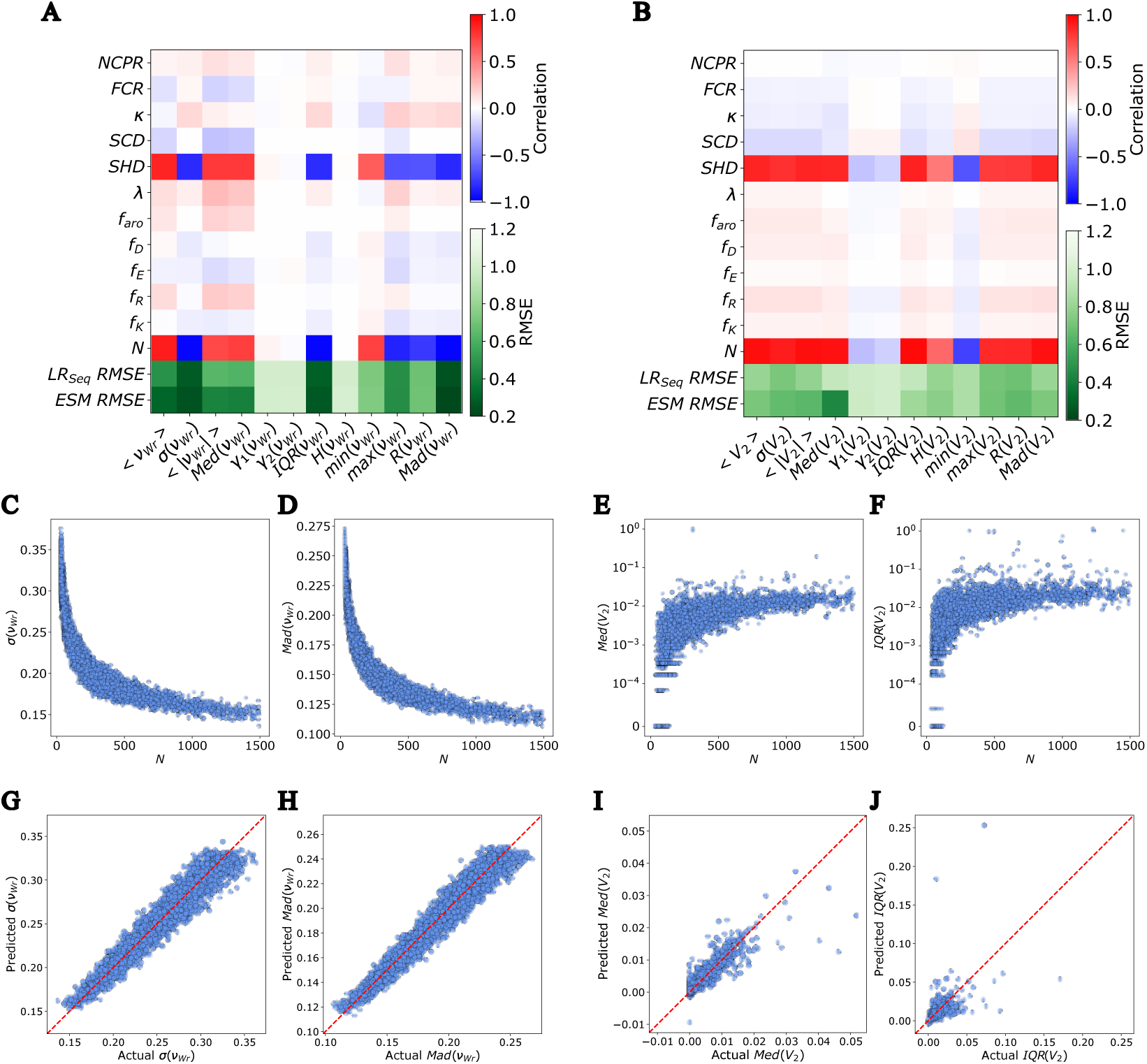
Sequence correlates of entanglement metrics. (A) Correlation matrix between sequence features and 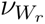 distribution features, and (B) correlation matrix for *V*_2_ distribution features. (C,D) Representative 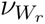 -based statistics, standard deviation (*σ*) and mean absolute deviation (*Mad*), shown as a function of chain length *N*, illustrating systematic size dependence across the database. (E,F) Representative *V*_2_-based statistics, median (*Med*) and interquartile range (*IQR*), shown as a function of *N*. (G,H) Predicted versus actual values for representative 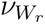 statistics using ESM-based neural network models. (I,J) Predicted versus actual values for representative *V*_2_ statistics using the same ESM-based models.

Across all sequence features, chain length emerged as the dominant correlate for both 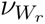 - and *V*_2_-based metrics (Fig. 2A and B, and Figs. S2 and S3). This trend is consistent with polymer physics: longer chains possess more configurational degrees of freedom and therefore greater capacity to accumulate coiling and higher-order entanglement. Among the sequence-derived descriptors, the strongest and most consistent correlations were observed for *SHD*, which quantifies hydrophobic residue patterning along the chain (Fig. 2C). Because *SHD* itself exhibits some dependence on chain length, these correlations should not be interpreted as strictly size-independent. Nevertheless, sequences with larger *SHD* values tend to adopt more cohesive conformations, increasing the frequency of intrachain contacts and thereby enhancing overall chain entanglement, as reflected in both 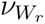 and *V*_2_. In contrast, the analogous charge patterning metric *SCD* shows weak correlations with either 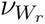 or *V*_2_. One possible explanation is that electrostatic interactions primarily modulate global conformational dimensions, whereas *V*_2_ reflects higher-order geometric organization that is less directly coupled to charge patterning.

Interestingly, these same compaction effects influence different aspects of the 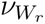 and *V*_2_ distributions in opposite ways. The standard deviation of 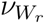, *σ*(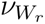), anticorrelates with *SHD*: sequences with strong hydrophobic patterning collapse into smoother, more uniformly curved conformations, which reduces frame-to-frame writhe variability. In other words, compaction damps the geometric fluctuations of the local coil structure. By contrast, *σ*(*V*_2_) correlates positively with *SHD*. Higher-order entanglement events require segments of the chain to approach and thread around one another, which becomes more frequent in compact states. These threading-like configurations are heterogeneous, giving rise to larger variability in *V*_2_ even as the overall chain becomes more cohesive. These observations indicate that hydrophobic patterning plays a disproportionate role in modulating the topological landscape of disordered chains, exerting distinct effects on local coiling versus higher-order entanglement.

To quantify the extent to which topological statistics are recoverable from sequence alone, we trained linear regression models using the full set of sequence features as predictors (Fig. 2A,B). Despite the simplicity of this approach, the models capture a substantial fraction of the variance for 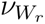 -based descriptors, yielding relatively low prediction errors quantified by root mean squared errors (RMSE) for distribution features such as the mean, standard deviation, *IQR*, and *Mad*. This indicates that writhe is largely determined by coarse-grained sequence features that modulate global chain dimensions and overall compaction, which are accessible to linear models. In contrast, the *V*_2_ descriptors are markedly more difficult to predict, and linear regression performs poorly for most of their distributional features. This disparity suggests that *V*_2_ reflects more intricate aspects of global topological organization that are not reducible to simple composition or patterning metrics.

To establish an approximate upper bound on predictability, we repeated the analysis using Evolutionary Scale Modeling (ESM) embeddings^48^ as inputs to an artificial neural network. These embeddings encode nonlinear and long-range sequence context learned from large-scale protein sequence data. Representative prediction plots are shown in Fig. 2G to J. As expected, ESM-based models substantially reduce the prediction error for both 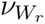 and *V*_2_ descriptors, with the most pronounced improvement observed for *V*_2_. Even with this richer representation, however, only the median of the *V*_2_ distribution are predicted with relatively low error, whereas higher-order moments remain difficult to capture.

Together, these analyses show that topological descriptors differ markedly in how readily their sequence dependence can be captured by current models. Writhe-based statistics are well predicted by both simple sequence features and ESM-based models, indicating that writhe reflects relatively low-order trends encoded in amino acid composition and patterning. In contrast, most *V*_2_-based statistics are poorly captured by linear models but improve with ESM embeddings, suggesting that higher-order entanglement is sequence-encoded yet depends on nonlinear and context-dependent features that are not accessible to simple composition or patterning metrics. The divergence in their predictability underscores the potential value of topological metrics for revealing distinct layers of sequence–structure relationships in IDPs.

### Structural correlates of entanglement metrics

An important question in evaluating the utility of topological descriptors is whether they capture information beyond standard structural measures used to characterize IDP ensembles. If 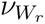 or *V*_2_ are fully determined by conventional observables such as the radius of gyration (*R_g_*) or pairwise distance statistics, then these metrics would largely recapitulate known global or local structural features. We therefore examined how the 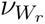 - and *V*_2_-based distribution features relate to a panel of structural properties (see Table S3) that quantify chain dimension, shape, and intra-chain contact formation.

Across the full IDRome database, 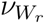 -based statistics show broad correlations with standard structural descriptors, including *R_g_*, end-to-end distance (*R*_ee_), hydrodynamic radius (*R_h_*), contact order, and shape metrics (Fig. 3A). Part of these correlations is expected because the global dimensions of an ensemble scale strongly with chain length, which also drives the overall increase of 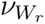 with *N*. More revealing is the relationship between 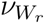 and the scaling exponent *ν*, which quantifies chain compactness independent of length. For IDPs, *ν* typically spans a range around the random-coil reference value of ∼ 0.5, with smaller values corresponding to more compact conformations and larger values to more expanded ensembles. 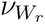 spans a comparable range but exhibits an opposite trend: sequences with smaller *ν* tend to display larger 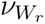 (Fig. S4). This anticorrelation suggests that enhanced compaction promotes increased local coiling, thereby amplifying geometric entanglement.

**Figure 3:**
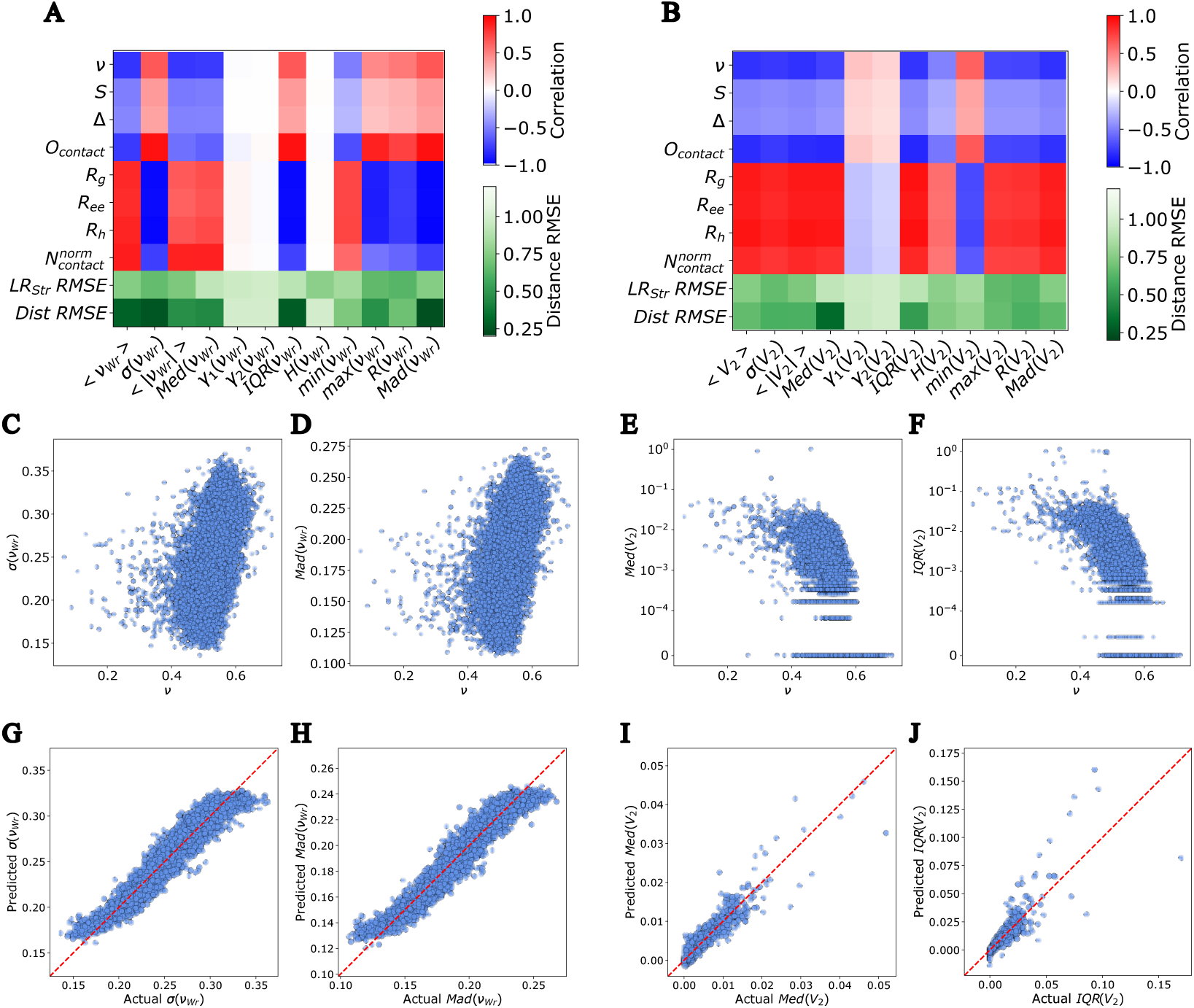
Structural correlates of entanglement metrics. (A,B) Correlation matrices between structural ensemble descriptors and distribution features of (A) 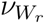 or (B) *V*_2_. (C,D) 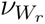 -based distribution features, including standard deviation (*σ*) and mean absolute deviation (*Mad*), shown as a function of the scaling exponent *ν*. (E,F) *V*_2_-based distribution features, including the median and interquartile range (*IQR*), shown as a function of *ν*. (G,H) Predicted versus actual values for representative writhe statistics using convolutional neural network models trained on pairwise distance matrices. (I,J) Predicted versus actual values for representative *V*_2_ statistics using the same modeling framework.

*V*_2_-based statistics also exhibit substantial correlations with multiple structural descriptors across the IDRome database (Fig. 3B), with several coefficients comparable in magnitude to those observed for 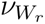 -based statistics. This indicates that higher-order entanglement features are not decoupled from global chain properties, but instead vary systematically with ensemble geometry. Notably, however, the pattern of correlations differs qualitatively from that of 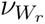 across multiple distribution features. In particular, measures of dispersion and extremal behavior, including the standard deviation, interquartile range, minimum, maximum, range, and mean absolute deviation, exhibit systematic sign reversals between 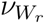 - and *V*_2_-based statistics. For both descriptors, the mean scaling indicates stronger entanglement in more compact chains with smaller *ν*. However, their fluctuations respond oppositely: increasing compactness amplifies the spread of 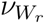 but reduces the spread of *V*_2_. This divergence implies that compaction enhances the heterogeneity of local coiling events contributing to 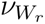, while simultaneously constraining the higher-order global entanglement captured by *V*_2_. Thus, 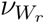 and *V*_2_ exhibit distinct fluctuation responses to conformational scaling despite sharing similar trends in their mean values.

To evaluate how much of the entanglement statistics can be inferred from structural information alone, we first trained linear regression models using ensemble-averaged structural descriptors as input features. These models show limited predictive power for both 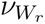 - and *V*_2_-based statistics (Fig. 3A,B), indicating that simple linear combinations of coarse structural descriptors are insufficient to recover the distributional properties of either topological measure. We therefore next trained a convolutional neural network (CNN) using full pairwise distance matrices as input, thereby leveraging the complete geometric information contained in the ensemble. This richer representation substantially improves predictive performance, particularly for 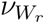 -based dispersion statistics such as the standard deviation and *Mad* (Fig. 3C, D, G and H). For *V*_2_, the CNN yields moderate accuracy for the median and *IQR* (Fig. 3E, F, I and J), but overall performance remains weaker and more scattered than for writhe. These results indicate that both topological descriptors require detailed geometric information beyond coarse ensemble averages. However, 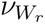 is more directly recoverable from pairwise distance structure, whereas *V*_2_ appears to depend on subtler spatial arrangements that are only partially captured even by full distance-matrix representations.

### Landscape of entanglement distribution features

Although the sequence- and structure-based analyses above identify which external features correlate with 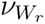 and *V*_2_, they do not reveal how the 12 distribution features for each descriptor relate to one another or which represent the dominant intrinsic sources of variation in the dataset. To examine the internal organization of these feature spaces directly, we applied t-distributed stochastic neighbor embedding (t-SNE)^49^ to all the 24 distribution features, embedding them into a two-dimensional manifold (Fig. 4A,B). This unsupervised embedding allows us to assess how the distribution features co-vary and whether a small number of low-dimensional collective variables capture most of the variation across the database.

**Figure 4:**
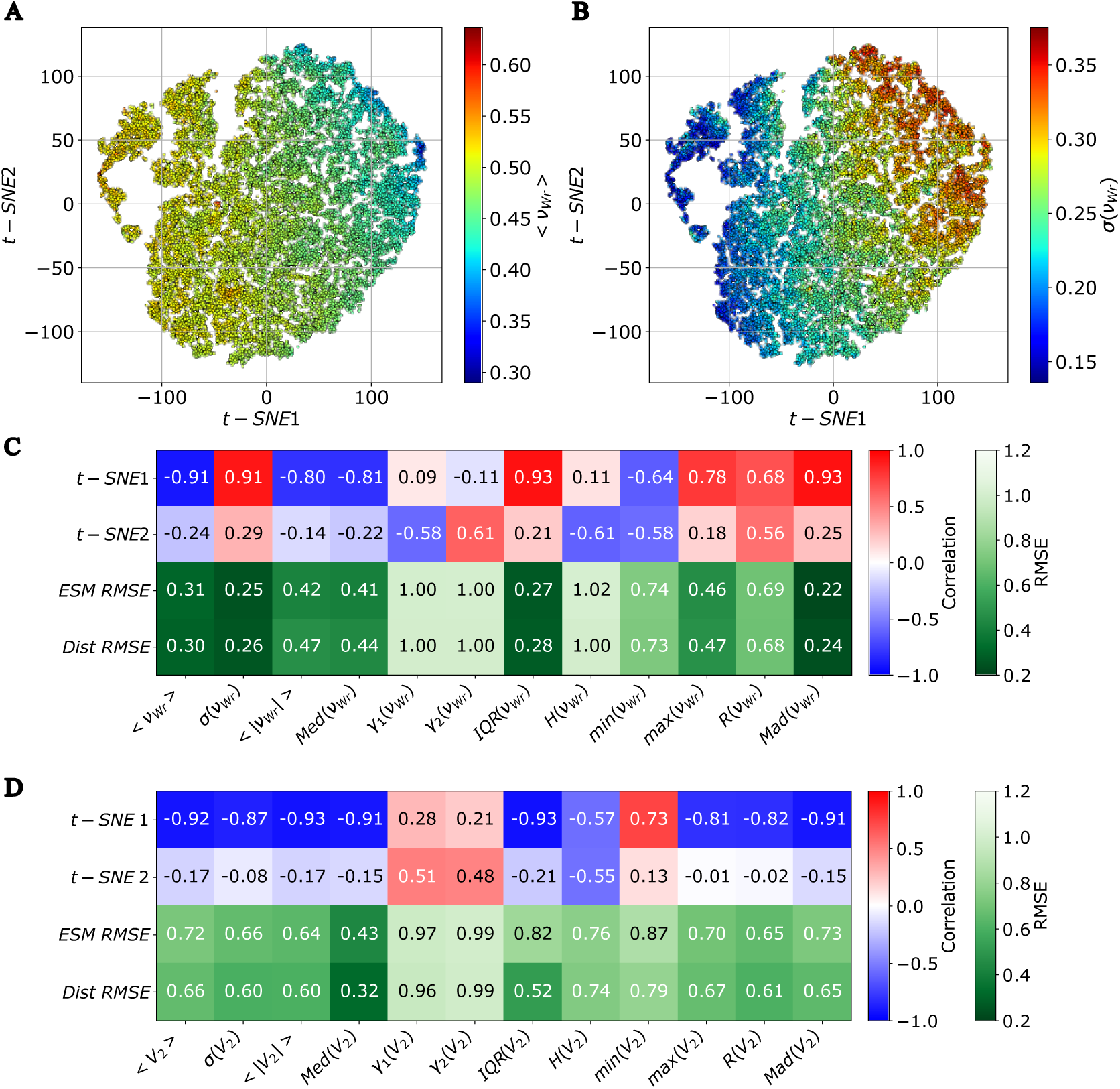
Landscape of 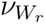 and *V*_2_ distribution features across the IDRome dataset. (A, B) Two-dimensional t-SNE embeddings constructed from the 24 distribution features for 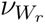 and *V*_2_, colored by the mean (A) and standard deviation (B) of the 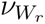. (C, D) Correlation matrices comparing each distribution feature with the first two t-SNE coordinates, in contrast to the RMSE of sequence-based (ESM) and structure-based (pairwise distance CNN) prediction models for each descriptor.

The t-SNE embeddings reveal coherent low-dimensional structure for both 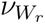 - and *V*_2_-based distribution features. For 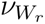 (Fig. 4C), the first embedding dimension exhibits strong correlations with multiple features, including the mean, standard deviation, median, *IQR*, range, and *Mad*, indicating that these features vary coordinately along a shared dominant dimension. This coordinated behavior suggests that much of the variability in 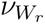 across IDRome is governed by a single underlying mode of geometric organization. A similarly strong alignment is observed for *V*_2_ (Fig. 4D and Fig. S6), where the first t-SNE dimension again correlates strongly with the mean and several dispersion-related features. Thus, despite differences in structural predictability, both descriptors exhibit internally coherent organization, with a principal low-dimensional axis, namely the first t-SNE coordinate (t-SNE 1), capturing a substantial fraction of their distributional variation.

The structure uncovered by t-SNE closely parallels the supervised learning results. Distribution features that align strongly with the dominant embedding axis tend to exhibit lower RMSE in both the ESM-based sequence model and the distance-matrix CNN model (Fig. 4C,D), whereas features weakly aligned with the embedding dimensions are more difficult to predict. This agreement indicates that the principal modes identified by unsupervised learning correspond to geometrically meaningful collective variables that are partially encoded in sequence and structural representations. Together, these results demonstrate that both 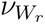 and *V*_2_ possess well-defined low-dimensional structure within the IDRome dataset, providing a compact and interpretable representation of topological variation that facilitates systematic comparison with functional annotations.

### Functional relevance of entanglement metrics

As shown in the previous section, the t-SNE embedding of the full set of entanglement distribution features revealed a small island of proteins with high values of 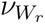 and *V*_2_, suggesting that these proteins share distinct topological characteristics. This observation raised the question of whether specific molecular functions are preferentially associated with particular regions of entanglement space, rather than being uniformly distributed across the IDRome dataset.

To examine potential functional structure, we performed a local Gene Ontology (GO) enrichment analysis across the t-SNE map using the Molecular Function terms. ^50^ GO enrichment analysis evaluates whether specific functional annotations are statistically overrepresented within a defined group of proteins compared to a background set. In this context, “local” refers to evaluating enrichment within neighborhoods defined by proximity in the low-dimensional entanglement embedding rather than across the entire dataset (see Methods). For each protein, enrichment within its neighborhood in entanglement space was summarized by a single score defined as the negative natural logarithm of the mean of the three lowest p-values. This score increases with the strength of functional enrichment and was used to construct a quantitative enrichment landscape over the embedding. As shown in Fig. 5A, this analysis revealed two prominent enriched regions located near the opposite extrema of the first t-SNE coordinate (t-SNE 1), indicating that specific molecular functions preferentially coincide with distinct entanglement signatures.

**Figure 5:**
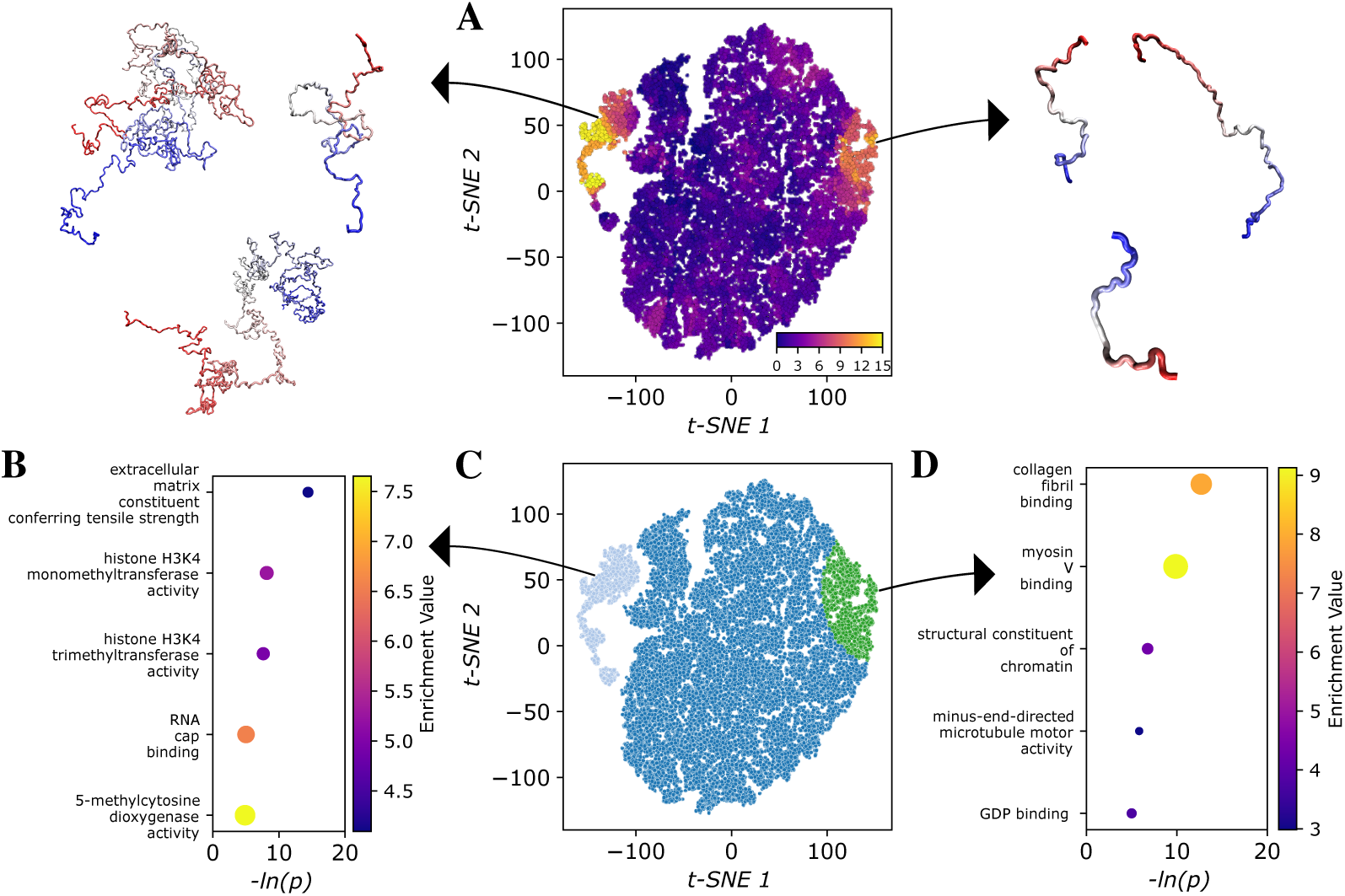
Functional relevance of entanglement metrics. (A) T-SNE embedding of all proteins colored by the strength of Gene Ontology (GO) enrichment analysis from each protein’s local neighborhood, quantified as the negative log of three lowest p-values. Representative simulated structures illustrating local entanglement at the two extrema of the first t-SNE coordinate are shown on either side of the embedding. (B) Top five enriched GO terms for the high-entanglement cluster. (C) Partition of the t-SNE map into two enriched clusters: a high-entanglement cluster (light blue) and a low-entanglement cluster (green). (D) Top five enriched GO terms for the low-entanglement cluster.

We further delineated these enriched neighborhoods into two regions, labeled Cluster 1 (light blue) and Cluster 2 (green), corresponding broadly to proteins with high and low entanglement characteristics, respectively (Fig. 5C, see Methods). Using the same GO enrichment framework described above, we then evaluated functional enrichment within each cluster and ranked the GO terms by fold enrichment magnitude and statistical significance to identify the dominant annotations. Cluster 1 was enriched in functions associated with “extracellular matrix constituent conferring tensile strength” and “histone H3K4 monomethyltransferase activity”. Proteins involved in extracellular matrix mechanics often operate under mechanical stress or participate in multivalent scaffold-like assemblies, where enhanced geometric entanglement may promote intermolecular coupling and mechanical resilience.^51^ Similarly, chromatin-associated factors involved in histone modification frequently engage in multivalent interactions within densely packed nuclear environments, where increased local chain entanglement may facilitate stable yet dynamic interaction networks.^52^ Cluster 2, in contrast, was enriched in “collagen fibril binding” and “myosin V binding”. These functions typically involve more directional or surface-specific interactions, where lower geometric entanglement may preserve accessibility of binding motifs and allow efficient recognition of structured partners. Although the two clusters exhibit distinct dominant annotations, partial overlap in extracellular matrix functions suggests that related biochemical roles can emerge from different underlying entanglement modes.

To assess the robustness of these findings and disentangle the contributions of individual entanglement descriptors, we repeated the t-SNE embedding and local GO enrichment analyses using either 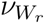 or *V*_2_ distribution features alone. The embedding constructed from *V*_2_ features recapitulated a separated group of proteins analogous to Cluster 1 and displayed similar top enriched Gene Ontology terms (Fig. S7), whereas the embedding based solely on 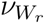 features did not produce a comparable isolated group or functional enrichment pattern (Fig. S8). Conversely, the 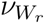 -only embedding reproduced enrichment patterns resembling those of Cluster 2, particularly in the low-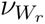 regime, while the *V*_2_-only embedding did not generate a corresponding hotspot. Together, these comparisons indicate that Cluster 1 is primarily driven by variation in *V*_2_, Cluster 2 by variation in 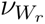, and that the two entanglement descriptors contribute complementary information to functional variation across the IDRome dataset.

Altogether, the emergence of two functionally enriched regions in entanglement space indicates that topological organization in IDP ensembles is not merely a geometric byproduct of chain flexibility but is structured in a manner that aligns with biological function. The differential contributions of 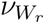 and *V*_2_ further suggest that distinct modes of chain entanglement underlie different classes of molecular activity, reinforcing the view that topological descriptors provide functionally relevant information beyond conventional structural metrics.

### Evolutionary aspects of entanglement metrics

The analyses above revealed that specific biological functions preferentially occupy distinct regions of entanglement space, suggesting that particular 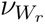 and *V*_2_ signatures may be required for proper molecular activity. This raises the question of whether such topological features are conserved across evolution. If entanglement contributes to functional constraints, orthologs of a given human protein should retain similar positions in entanglement space despite substantial sequence divergence.

To investigate the evolutionary conservation of these entanglement signatures, we selected representative human proteins from the two enriched clusters and from a background region located in the middle of the t-SNE embedding (Fig. 6A). For each enriched cluster, we chose proteins corresponding to the top two GO terms identified in the previous section. For the central background region, we selected six proteins associated with GO terms that were present but not strongly enriched or spatially localized to either hotspot, thereby providing a baseline comparison. The resulting set of human sequences, along with their residue ranges and the number of available orthologs, is listed in Supporting Table S4, which includes all UniProt identifiers used in the analysis. These proteins served as the basis for an evolutionary survey of entanglement conservation.

**Figure 6:**
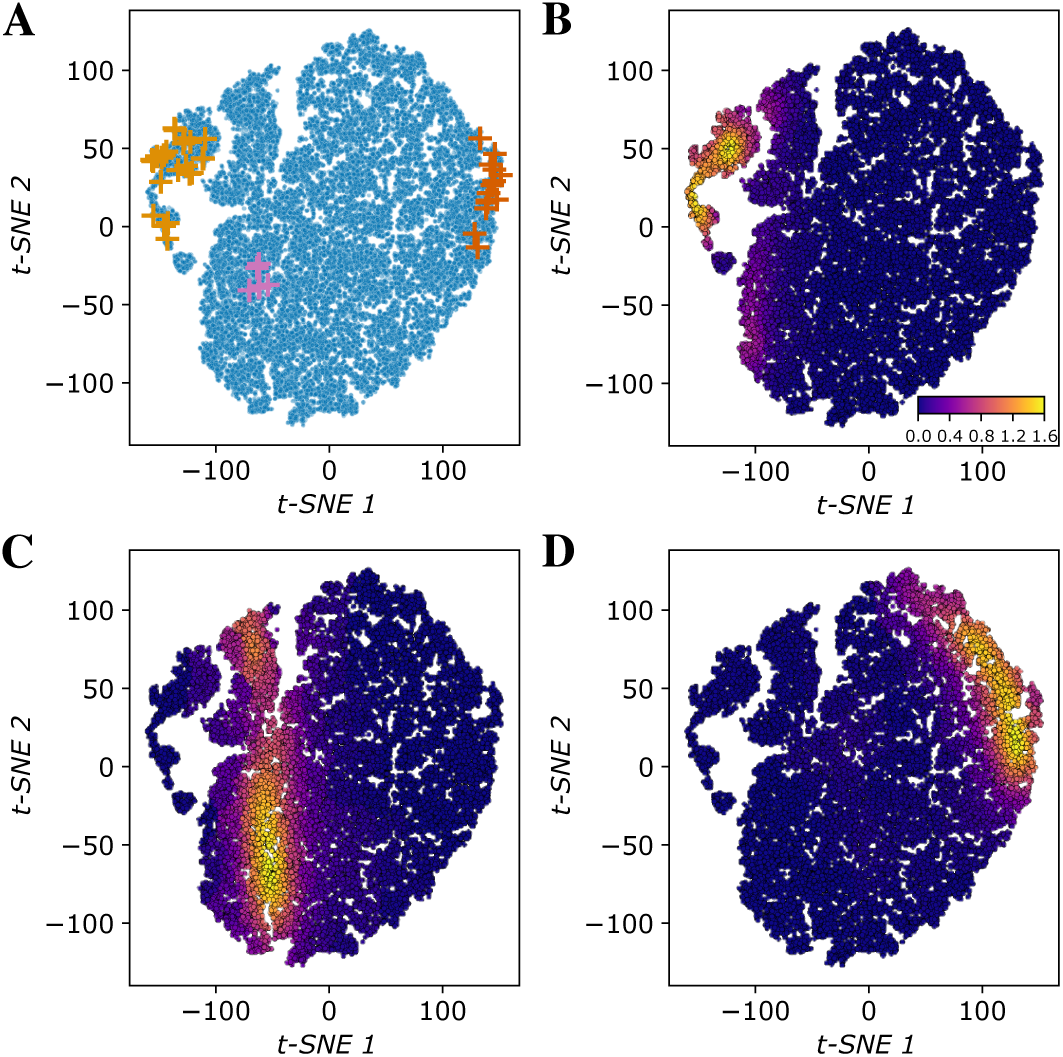
Conservation of entanglement metrics across orthologs. (A) Locations of the selected human IDP sequences used for ortholog analysis plotted on the entanglement t-SNE embedding. Cluster 1 (light orange), Cluster 2 (dark orange), and Cluster 3 representing the background sequences (pink) are highlighted. (B–D) Density projections of ortholog entanglement signatures onto the same t-SNE embedding for Cluster 1 (B), background sequences (C), and Cluster 2 (D).

We then acquired orthologs from the Ensembl database for the proteins selected from Cluster 1, Cluster 2, and Cluster 3 (the background region). Each ortholog was simulated using coarse-grained molecular dynamics, generating a total of 1.5 *µ*s of production trajectories per sequence. Using these simulations, we calculated the 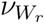 and *V*_2_ values, yielding ensemble-level topological descriptors directly comparable to those of the corresponding human proteins.

To determine whether orthologs share similar entanglement signatures with their human counterparts, we projected ortholog sequences onto the human entanglement t-SNE space. For each ortholog, we computed its Euclidean distance to all human entanglement distribution vectors, identified the 100 nearest neighbors in the full entanglement distribution feature space, and calculated a weighted average of their t-SNE coordinates using an exponential kernel. These projected coordinates were used to construct kernel density estimates over the human t-SNE map, yielding density distributions for orthologs derived from Cluster 1, Cluster 2, and the background set (Fig. 6B to D).

The resulting projections reveal clear patterns of conservation across species. Orthologs of Cluster 1 proteins strongly localize to the same region as their human counterparts, indicating preservation of high-entanglement signatures. Cluster 2 orthologs likewise occupy similar regions of entanglement space, although with greater dispersion along the second SNE coordinate compared to Cluster 1. In contrast, orthologs from the background set display broader spreading across the embedding, particularly along t-SNE 2. Notably, however, background orthologs remain relatively aligned along t-SNE 1, suggesting that certain aspects of their entanglement regime are still maintained despite weaker hotspot enrichment. Together, these results show that proteins within functionally enriched entanglement clusters retain their topological signatures more tightly across evolution than proteins without enriched functions.

We further examined whether evolutionary divergence between species correlates with changes in entanglement features. For each ortholog–human pair, we compared the phylogenetic divergence time with the corresponding entanglement distance in the t-SNE embedding space. If entanglement signatures systematically vary over evolutionary time, one would expect a positive/negative correlation between divergence time and entanglement distance. Across all proteins analyzed, the correlations were weak in magnitude, ranging from Pearson coefficients of approximately −0.3 to 0.2, with no consistent directional trend (Fig. S9). This lack of systematic correlation indicates that entanglement distances do not progressively increase or decrease with evolutionary separation. Together, these results suggest that entanglement features remain relatively stable across substantial evolutionary timescales, consistent with the presence of functional constraints on topological organization.

## Conclusion

This work establishes entanglement as a meaningful and previously underexplored dimension for characterizing the conformational ensembles of intrinsically disordered proteins (IDPs). While IDP ensembles are traditionally described using measures of size, shape, or pairwise contact formation, such descriptors do not fully capture how a chain collectively winds and organizes in three-dimensional space. By introducing distribution-level representations of the writhe scaling exponent 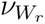 and the higher-order invariant *V*_2_ from mathematical knot theory, we show that entanglement provides a compact and interpretable axis of ensemble variation across the IDRome database.^44^ The resulting entanglement landscape is structured rather than random, revealing coherent low-dimensional organization that integrates geometric information across scales.

The two entanglement descriptors examined here capture complementary aspects of chain organization. 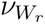 primarily reflects systematic patterns of local coiling that scale with chain compaction and are largely recoverable from coarse sequence features and established structural descriptors. In contrast, *V*_2_ detects higher-order global topological complexity that depends on coordinated multi-segment configurations and exhibits weaker predictability from simple sequence composition or known structural metrics. These distinctions are evident in their opposite fluctuation responses to compaction and in their differential recoverability from sequence- and structure-based models, indicating that entanglement cannot be captured by a single statistic, but instead reflects distinct and partially independent aspects of chain organization.

Most importantly, entanglement aligns with biological function and evolutionary constraint. Proteins cluster into distinct regions of entanglement space enriched for specific molecular activities, and these signatures are preferentially retained across orthologs for proteins drawn from enriched clusters. At the same time, the absence of systematic drift in entanglement distance with evolutionary divergence indicates that these features remain relatively stable over evolutionary timescales. Together, these findings indicate that entanglement is not merely a geometric byproduct of disorder but a biologically relevant organizational feature that helps connect sequence, ensemble structure, and molecular function in IDPs.

## Method

### Writhe and *V*_2_ calculation

For each conformation in an IDP ensemble, we computed the geometric writhe (*Wr*) and the second Vassiliev invariant (*V*_2_) following standard definitions for polygonal chains. Both *Wr* and *V*_2_ are continuous functions of the protein’s coordinates in the three-dimensional space.

The writhe of a curve in three-dimensional space quantifies the degree to which the curve winds around itself. It is defined as the Gauss linking integral over one curve: ^53^

#### Definition 0.1

*(Writhe) For an oriented curve l with an arc-length parametrization γ, the writhe, Wr, is the double integral over l:*

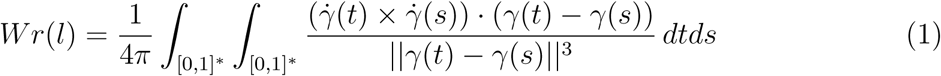

*where γ̇ denotes the derivative of γ, and the integral runs over* [0, 1]^∗^×[0, 1]^∗^ *for all s, t* ∈ [0, 1] *such that s* ≠ *t*.

The writhe can have both positive and negative values depending on the orientation of a curve. Even though a high absolute writhe value may indicate geometric complexity, the writhe is sensitive to local geometrical entanglement. It can also be expressed as the average algebraic sum of signs of all crossings in a projection of a curve with itself over all possible projection directions. For polygonal curves, it can be expressed as a finite sum of signed geometric probabilities that any two edges cross in any projection direction, which can be computed exactly.^54^

*V*_2_ is a higher-order topological invariant that detects more complex entanglement features, including near-threading events.^40^

#### Definition 0.2

*Let j*_1_ *< j*_2_ *< j*_3_ *< j*_4_ *be points on a knot or link diagram. We will say that this four-tuple corresponds to an alternating crossing when j*_1_*, j*_3_ *and j*_2_*, j*_4_ *are two crossing points in the diagram such that if j*_1_ *belongs to the over-arc in the crossing, j*_2_ *belongs to the under-arc in the crossing or vice-versa*.

#### Definition 0.3 (Second Vassiliev measure)

*For an oriented curve l with the arc-length parametrization γ, the second Vassiliev measure, V*_2_*, is the double alternating self-linking integral over l:*

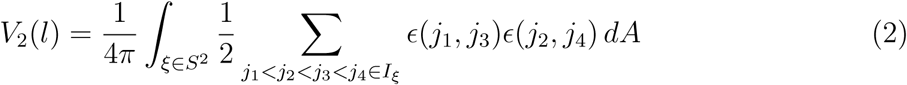

*where ɛ*(*s, t*) = ±1 *is the sign of the crossing between the projection of γ*(*s*) *and γ*(*t*)*, and where I_ξ_ denotes the set of pairs of alternating crossings in the projection to the plane with normal vector ξ*.

In practice, for each sequence in the IDRome dataset, writhe was evaluated for all 1000 frames of the simulation trajectory, yielding the full frame-by-frame writhe distribution for every ensemble. The computation of *V*_2_ is substantially more expensive, as it requires averaging over projection directions of the planar crossing diagram for each frame. For an open polygonal chain, *V*_2_ was computed as the average over multiple projection directions, and in this work we used 3000 uniformly sampled projections per frame. To maintain computational tractability, *V*_2_ was evaluated on 200 evenly spaced frames per sequence, which uniformly sample the full trajectory. Convergence with respect to both the number of projection directions and the number of sampled frames was verified (Fig. S1). The resulting set of *V*_2_ values defines the ensemble-level *V*_2_ distribution for each protein.

### Sequence- and structure-based prediction of entanglement metrics

To evaluate the extent to which entanglement descriptors can be inferred from either primary sequence or ensemble geometry, we developed two supervised learning models based on sequence embeddings and structural distance representations. For sequence-based prediction, each protein sequence was embedded using the 605M-parameter ESM-2 model^48^ yielding high-dimensional representations that capture contextual and evolutionary information. A feed-forward neural network was trained on these embeddings to regress the corresponding 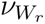- and *V*_2_-based distribution features. The network comprises three hidden layers with 256, 128, and 64 units, respectively, each followed by a SiLU activation function, with dropout (*p* = 0.1) applied after the first two layers, and a final linear output neuron for scalar prediction. Models were optimized by minimizing the mean squared error (MSE) loss using the AdamW optimizer^55^ (learning rate 3 × 10^−4^, weight decay 1 × 10^−4^), together with a ReduceLROnPlateau scheduler, gradient clipping (maximum norm 2.0), and early stopping (patience of 20 epochs). Data were split into training, validation, and test sets (approximately 70/15/15), and performance was evaluated using the root-mean-square error (RMSE) on the held-out test set.

For structure-based prediction, we computed the mean pairwise distance matrix across the conformational ensemble for each sequence and treated the resulting matrices as image-like inputs. A convolutional neural network (CNN)^56^ was then trained on these matrices to predict the same entanglement descriptors. Each input matrix was first normalized on a per-sample basis using min-max scaling to [0, 1] and resized to a fixed resolution of 64 × 64 via bilinear interpolation. The network consists of four convolutional blocks with channel dimensions 1 → 32 → 64 → 128 → 128, each using 3 × 3 kernels with padding and SiLU activations; max-pooling layers follow the first three convolutions, and a global adaptive average pooling layer reduces the feature map to a 128-dimensional vector. The regression head comprises two fully connected layers (128→64→1) with a SiLU activation between them to produce a single scalar output. Models were trained and evaluated with the same setups as the feed-forward neural network.

### Local Gene Ontology enrichment analysis

To quantify functional enrichment across the entanglement embedding, we performed a local Gene Ontology (GO) analysis using Molecular Function annotations for each protein in the IDRome dataset. For a given protein, we defined a neighborhood consisting of 1900 proteins including itself, obtained by selecting its 1899 nearest neighbors in the full entanglement feature space. This neighborhood size approximates the number of sequences contained in the isolated island located at the positive extreme of the first t-SNE coordinate and ensures a consistent neighborhood size across all proteins.

Within each neighborhood, GO enrichment was evaluated using fold enrichment and Fisher’s exact test based on the presence or absence of each GO Molecular Function term among the neighboring proteins. Fold enrichment was defined as (*k/n*)*/*(*K/N*) where *k* is the number of proteins annotated with a given GO term within the neighborhood, *n* is the neighborhood size, *K* is the number of proteins annotated with that term in the full dataset (background), and *N* is the total number of proteins in the dataset.

Statistical significance was assessed using a one-sided Fisher’s exact test for overrepresentation, with p-value

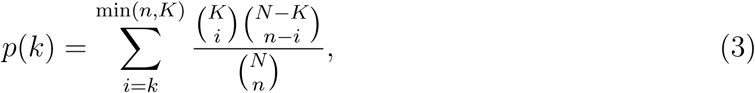

where variables are defined the same as in the fold enrichment equation. The p-values were further adjusted for multiple hypothesis testing using the Benjamini–Hochberg false discovery rate procedure.^57^

Only GO terms with fold enrichment greater than 3.5 were retained to reduce low-effect signals. For each protein, the three smallest adjusted p-values among the retained terms were identified, their mean was computed, and the negative natural logarithm of this mean was defined as the local enrichment score. These scores were mapped onto the t-SNE embedding to construct the enrichment landscape shown in Fig. 5A. Sensitivity analyses over neighborhood sizes ranging from 1500 to 2500 proteins and enrichment thresholds from 3 to 4 produced qualitatively similar hotspot structures.

To further characterize the two prominent enrichment hotspots observed in Fig. 5A, we performed cluster-level GO enrichment analyses. Cluster 1 was defined to isolate the small island located at the negative extreme of the first t-SNE coordinate. Cluster 2 was defined by selecting a representative protein from the positive extreme of the first t-SNE coordinate and constructing a cluster of 2000 proteins based on Euclidean distances in the t-SNE space. GO enrichment within each cluster was evaluated using the same fold enrichment and Fisher’s exact testing framework described above. GO terms were ranked by fold enrichment and statistical significance. The top five terms were selected from those with fold enrichment among the top 40 values and adjusted p-values below the mean of the smallest 40 adjusted p-values, and were further ranked according to their adjusted p-values.

### Ortholog simulations and projections

Orthologs were selected for representative human proteins drawn from the two most enriched GO terms within each of the two entanglement clusters, as well as six additional proteins from the central region of the t-SNE embedding serving as a background set. Ortholog sequences were retrieved from the Ensemble database,^58^ and the intrinsically disordered regions (IDRs) were defined by mapping the annotated human IDR boundaries onto the ortholog sequence alignment. All ortholog IDRs were simulated using the CALVADOS2 model^59^ implemented in OpenMM.^60^ All simulations were run using a Langevin thermostat at 310 K with a friction coefficient of 0.01 ps^−1^ and a time step of 10 fs. A total of 1.5 µs simulation was performed for each sequence, with the first 0.5 µs discarded as equilibration. From the production trajectories, 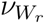 values were computed from 1000 equally spaced frames per sequence, while *V*_2_ values were computed from 200 equally spaced frames per sequence.

To compare ortholog entanglement signatures with those of human proteins, orthologs were projected onto the previously obtained human t-SNE embedding. For each ortholog, Euclidean distances were computed between its entanglement distribution feature vector and those of all human proteins. The 100 nearest human neighbors in entanglement feature space were identified, and their corresponding t-SNE coordinates were used to estimate the ortholog position via a weighted average. Weights were assigned using a Gaussian kernel of the form 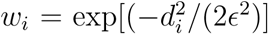, where *d_i_* is the Euclidean distance in entanglement feature space and *ɛ* was set to the mean distance between each human protein and its 100 nearest neighbors. The weighted average of the 100 nearest human t-SNE coordinates defined the projected ortholog position. Kernel density estimation was then applied to the projected ortholog positions in the two-dimensional t-SNE space to obtain the density distributions shown in Fig. 6.

To quantify evolutionary conservation beyond spatial localization in the embedding, divergence times (in unit of MYA, millions of years ago) between each human protein and its orthologs were obtained using the TimeTree database. ^61^ For each ortholog–human pair, we computed the Euclidean distance between their projected t-SNE coordinates and calculated the Pearson correlation coefficient between divergence time and entanglement distance.

## Supporting Information Available

Supporting tables for all sequence, structure and distribution features, and figures illustrating additional parameters.

## Acknowledgement

This work is supported by the National Institutes of Health (R35GM146814, W.Z.) and (R01GM152735, E.P.). The authors also acknowledge support from the SCENE program (H.S.) and from Research Computing at Arizona State University.

## Notes

### Competing Interest Statement

The authors have declared no competing interest.

