## Supplementary figures and tables. for "Topological Entanglement in Intrinsically Disordered Proteins: Sequence, Structural, and Functional Determinants"

### Supporting Information for: "Topological Entanglement in Intrinsically Disordered Proteins: Sequence, Structural, and Functional Determinants"

Wangfei Yang[1*], Henry Silvernail[2*], Debasis Saha[1],Eleni Panagiotou[3], and Wenwei Zheng[1,4a]

[1] College of Integrative Sciences and Arts, Arizona State University, Mesa, AZ 85212, USA

[2] Brophy College Preparatory, Phoenix, AZ 85012, USA

[3] School of Mathematical and Statistical Sciences, Arizona State University, Tempe, AZ 85282, USA

[4] Center for Biological Physics, Arizona State University, Tempe, AZ 85282, USA

∗ These authors contributed equally to this work.

[a]Electronic mail:

### 1 Supplementary Figures

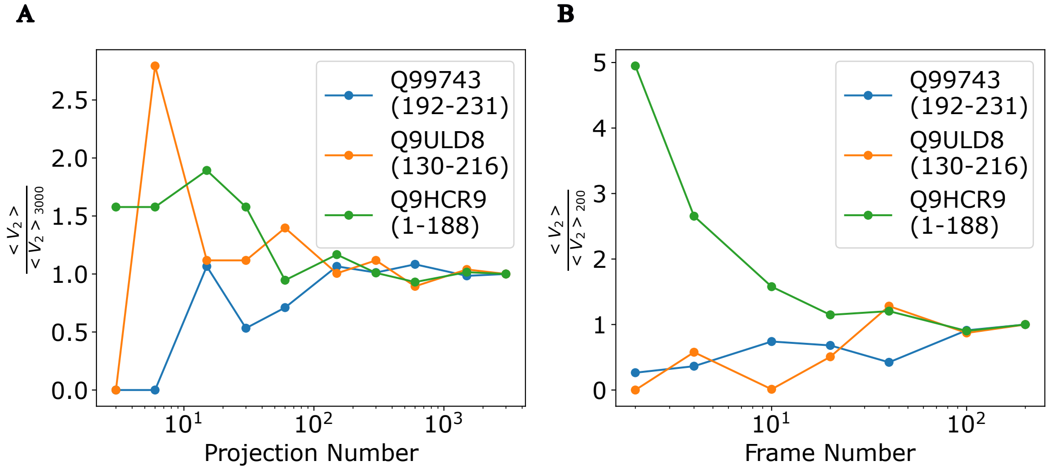


Figure S1: **Convergence of $V_2$ with respect to the number of projections and the number of frames.** (A) For a single frame in the trajectory, we show the ratio of the averaged $V_2$ computed using a given number of random projections to that computed using 3000 projections, which is taken as the converged reference value. (B) For an entire simulation trajectory, we show the ratio of the averaged $V_2$ computed using a given number of frames to that computed using 200 frames, which is taken as the converged reference value. The legend indicates representative sequences, including their UniProt identifiers and residue ranges corresponding to the disordered regions.

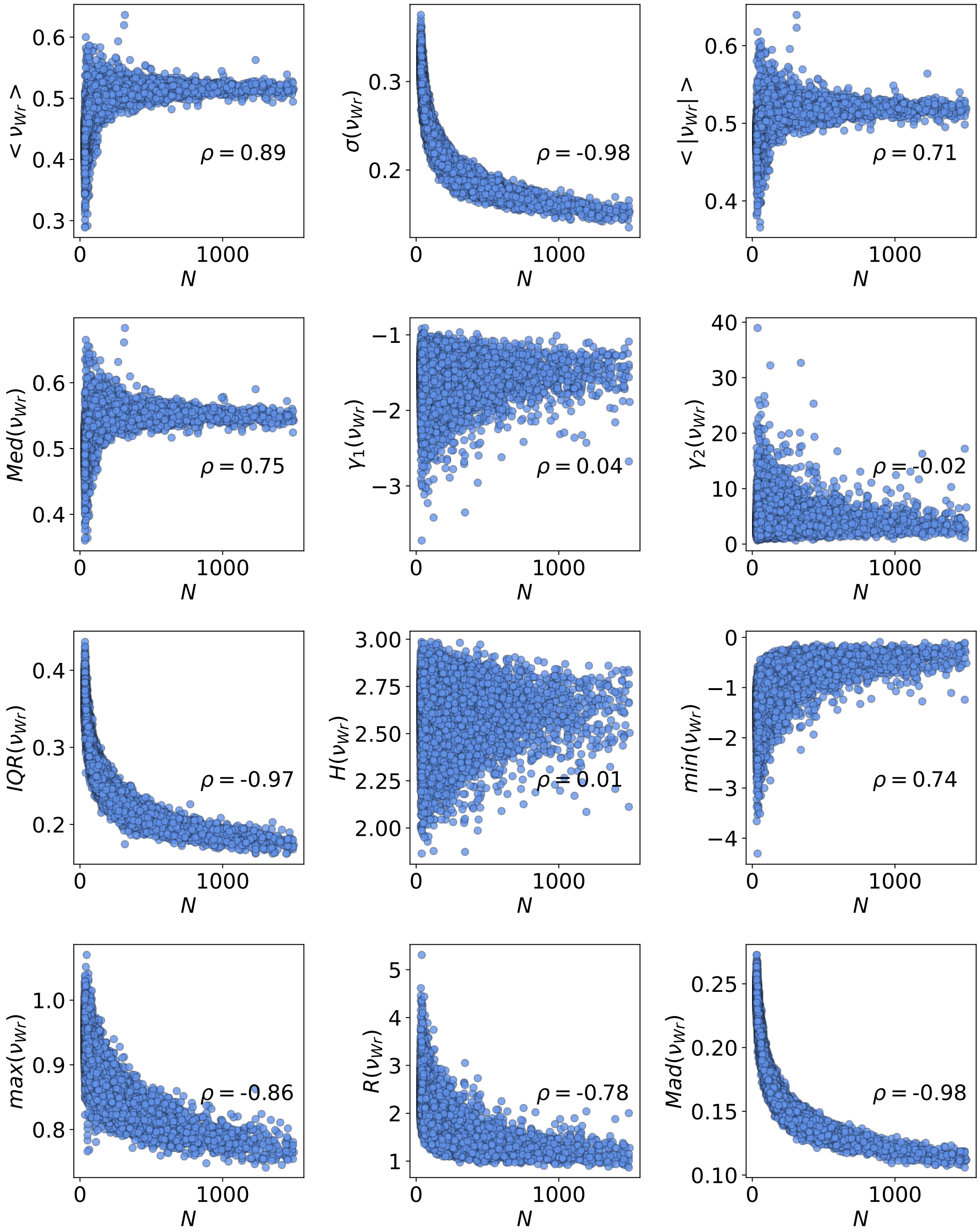


Figure S2: **Correlation plots between $\nu_{Wr}$ distribution features and the chain length $N$.**

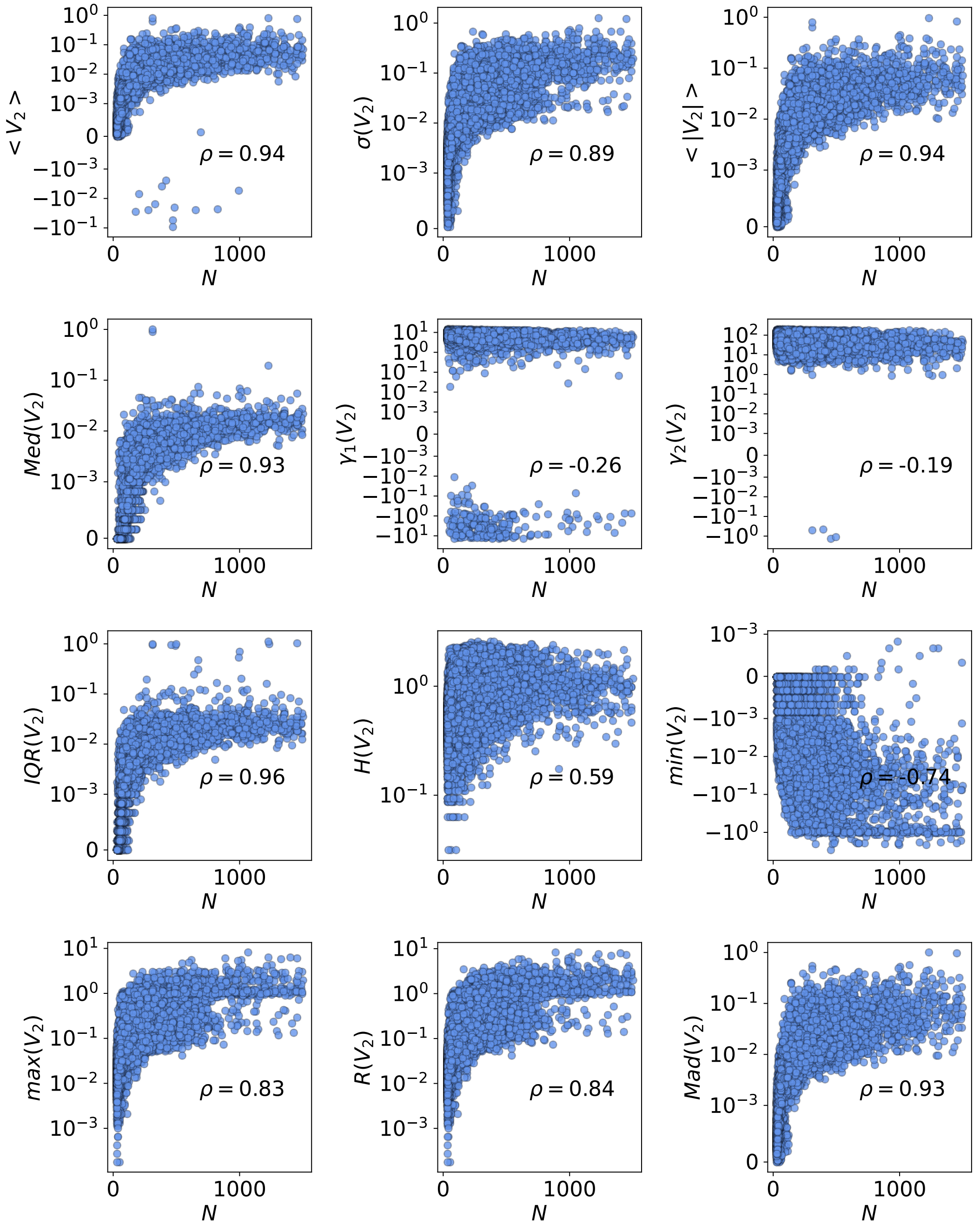


Figure S3: **Correlation plots between $V_2$ distribution features and the chain length $N$.**

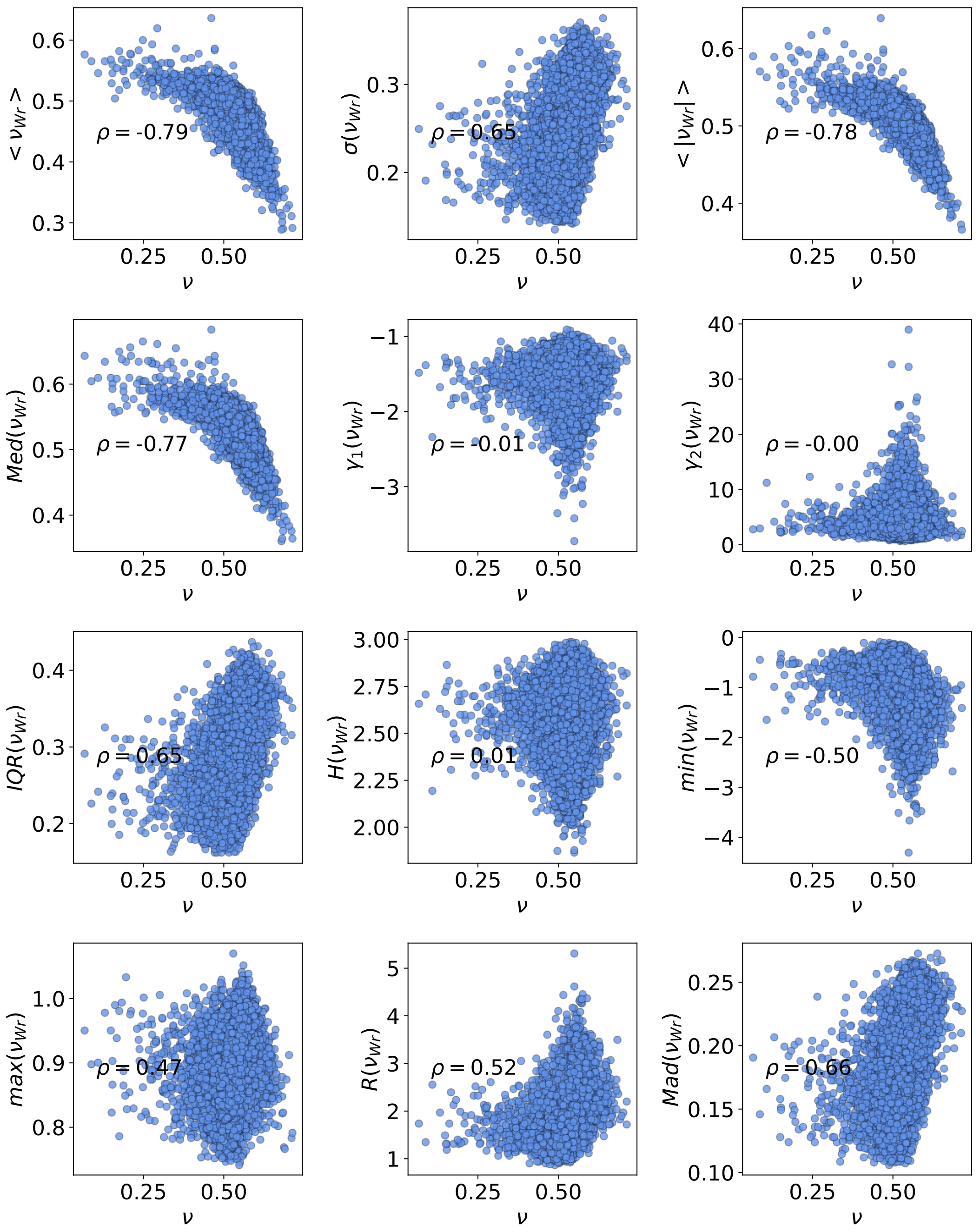


Figure S4: **Correlation plots between $\nu_{Wr}$ distribution features and the scaling exponent $\nu$.**

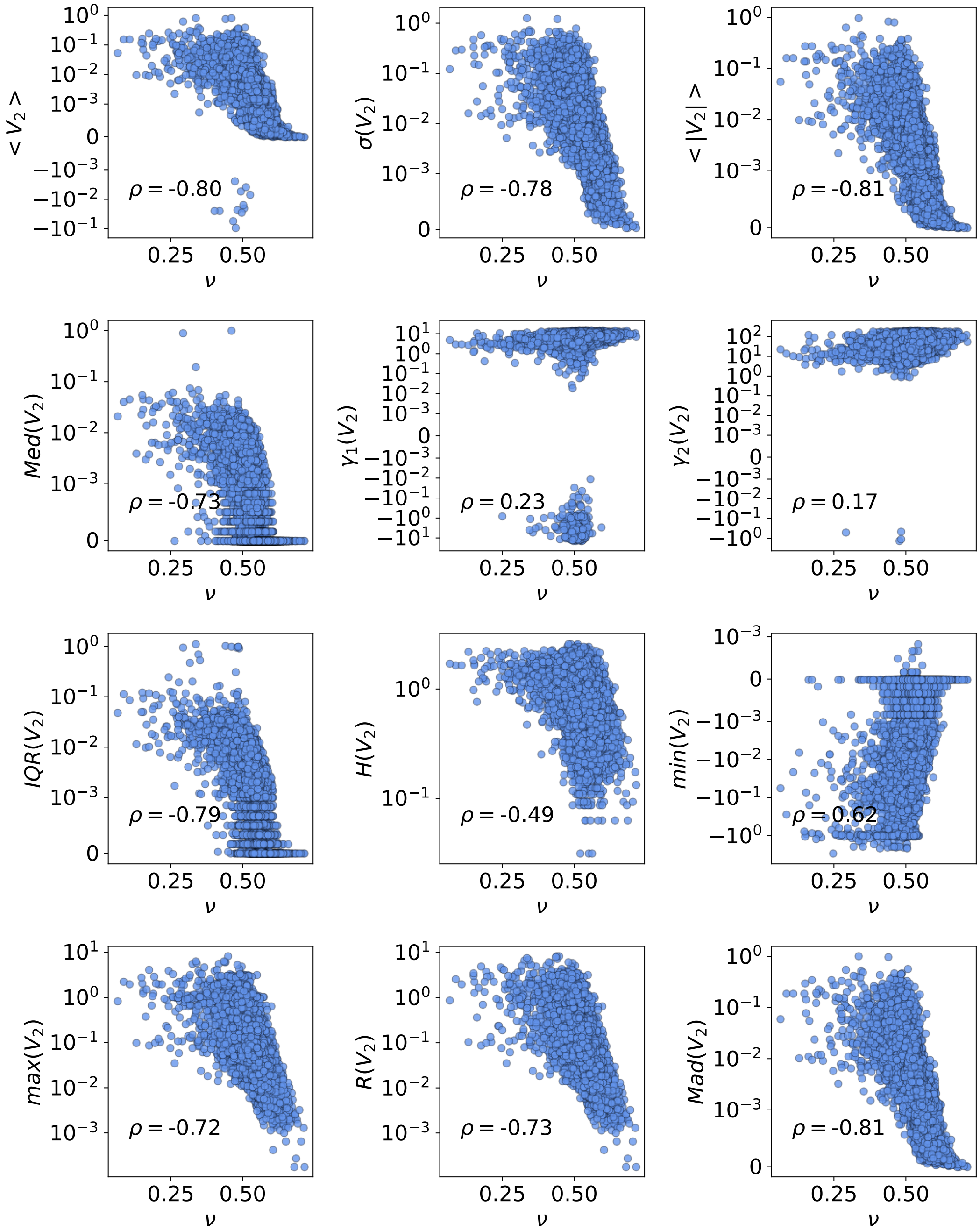


Figure S5: **Correlation plots between $V_2$ distribution features and the scaling exponent $\nu$.**

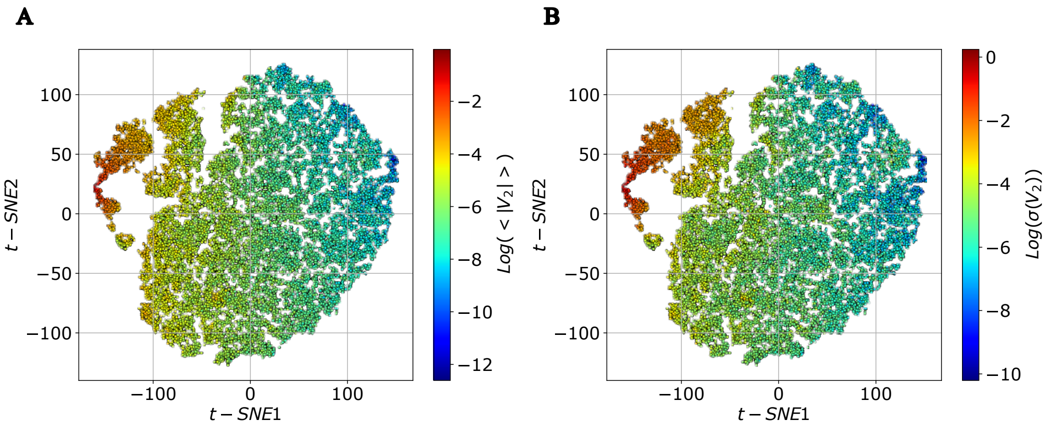


Figure S6: Two-dimensional t-SNE embeddings constructed from the 24 distribution features for $\nu_{W_r}$ and $V_2$, colored by the mean absolute (A) and standard deviation (B) of the $V_2$.

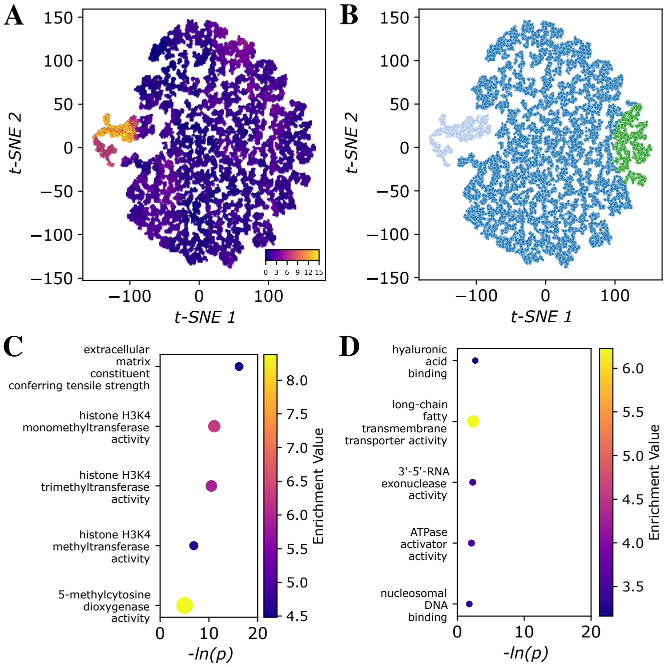


Figure S7: **Functional relevance of topological variation in the IDRome dataset using only $V_2$.** (A) T-SNE embedding of all proteins colored by the strength of Gene Ontology (GO) enrichment analysis from each protein's local neighborhood, quantified as the negative log of three lowest p-values. (B) Partition of the t-SNE map into two enriched clusters: a high-entanglement cluster (light blue) and a low-entanglement cluster (green). (C) Top five enriched GO terms for the high-entanglement cluster. (D) Top five enriched GO terms for the low-entanglement cluster.

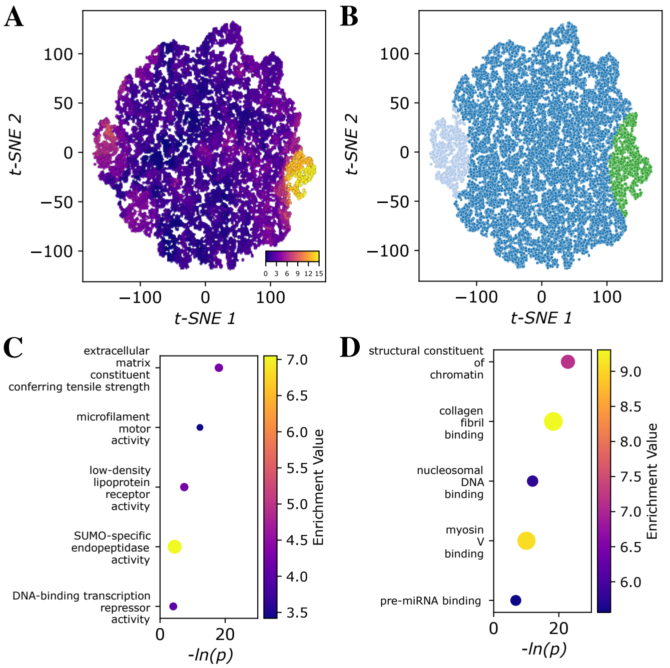


Figure S8: **Functional relevance of topological variation in the IDRome dataset using only $\nu_{W_r}$.** (A) T-SNE embedding of all proteins colored by the strength of Gene Ontology (GO) enrichment analysis from each protein's local neighborhood, quantified as the negative log of three lowest p-values. (B) Partition of the t-SNE map into two enriched clusters: a high-entanglement cluster (light blue) and a low-entanglement cluster (green). (C) Top five enriched GO terms for the high-entanglement cluster. (D) Top five enriched GO terms for the low-entanglement cluster.

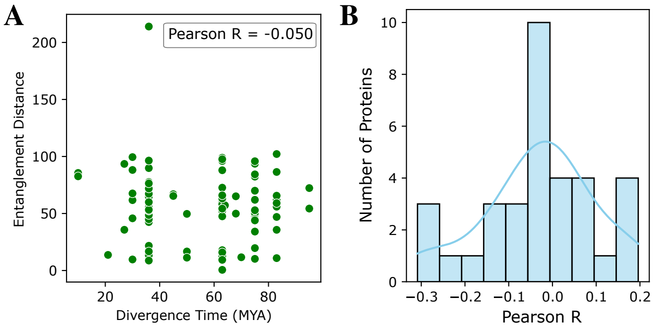


Figure S9: **Conservation of topological features across taxonomic distances.** (A) Relationship between the entanglement distance between each ortholog and its corresponding human protein and evolutionary divergence time (in unit of MYA, millions of years ago). The Pearson correlation coefficient indicates no significant dependence of topological distance on divergence time. (B) Distribution of Pearson correlation coefficients quantifying the relationship between evolutionary distance and entanglement distance across all proteins analyzed.

### 2 Supplementary Tables

Table S1: Sequence features. All sequence features were provided by the IDRome database[1].

| Name | Abbreviation | Definition / Equation |
|---|---|---|
| Net Charge Per Residue | NCPR | $\frac{N_+ - N_-}{N}$ |
| Fraction of Charged Residues | FCR | $\frac{N_+ + N_-}{N}$ |
| Charge Patterning Parameter | $\kappa$ | Das–Pappu charge segregation metric[2] |
| Sequence Charge Decoration | SCD | $\frac{1}{N}\sum_{i<j} q_i q_j \sqrt{j-i}$[3] |
| Sequence Hydropathy Decoration | SHD | $\frac{1}{N}\sum_{i<j} (\lambda_i \, \lambda_j) \, (j-i)^{-1}$[4] |
| Average Hydropathy | $\lambda$ | $\frac{1}{N}\sum_{i=1}^{N} \lambda_i$ |
| Aromatic Fraction | $f_{\mathrm{aro}}$ | $\frac{N_{F,W,Y}}{N}$ |
| Aspartate Fraction | $f_D$ | $\frac{N_D}{N}$ |
| Glutamate Fraction | $f_E$ | $\frac{N_E}{N}$ |
| Arginine Fraction | $f_R$ | $\frac{N_R}{N}$ |
| Lysine Fraction | $f_K$ | $\frac{N_K}{N}$ |
| Chain Length | $N$ | Total number of residues |

Table S2: Distribution features to characterize $\nu_{Wr}$ and $V_2$ ensembles

| **Name** | **Symbol** | **Equation** |
|---|---|---|
| Mean | $<>$ | $\mu = \frac{1}{N}\sum_{i=1}^{N} x_i$ |
| Standard deviation | $\sigma$ | $\sqrt{\frac{1}{N}\sum_{i=1}^{N}(x_i - \mu)^2}$ |
| Mean absolute value | $< \| \| >$ | $\frac{1}{N}\sum_{i=1}^{N} \|x_i\|$ |
| Median | $Med$ | median$(x_1, \ldots, x_N)$ |
| Skewness | $\gamma_1$ | $\frac{\frac{1}{N}\sum_{i=1}^{N}(x_i - \mu)^3}{\sigma^3}$ |
| Kurtosis (excess) | $\gamma_2$ | $\frac{\frac{1}{N}\sum_{i=1}^{N}(x_i - \mu)^4}{\sigma^4} - 3$ |
| Interquartile range | IQR | the difference between the 75th and 25th percentiles |
| Entropy | $H$ | $-\sum_{i} p_i \log p_i$ (where $p_i$ is the probability density of the $i$-th bin) |
| Minimum | min | $\min(x_i)$ |
| Maximum | max | $\max(x_i)$ |
| Range | R | $\max(x_i) - \min(x_i)$ |
| Mean absolute deviation | $Mad$ | $\frac{1}{N}\sum_{i=1}^{N} \|x_i - \mu\|$ |

Table S3: Structural features. All structural featrues other than contact order and normalized number of contacts were provided by the IDRome database[1].

| **Name** | **Symbol** | **Definition / Equation** |
|---|---|---|
| Scaling exponent | $\nu$ | Polymer scaling exponent from intra-chain distance fitting |
| Asphericity | $S$ | $\frac{(\lambda_1-\lambda_2)^2+(\lambda_2-\lambda_3)^2+(\lambda_3-\lambda_1)^2}{2(\lambda_1+\lambda_2+\lambda_3)^2}$ in which $\lambda_i$ are the eigenvalues of the gyration tensor |
| Shape anisotropy | $\Delta$ | $1-3\,\frac{\lambda_1\lambda_2+\lambda_2\lambda_3+\lambda_3\lambda_1}{(\lambda_1+\lambda_2+\lambda_3)^2}$ |
| Contact order | $O_{\text{contact}}$ | $\frac{1}{N_{\text{contact}}}\sum_{(i,j)\in\text{contacts}}\frac{\lvert i-j\rvert}{N}$ |
| Radius of gyration | $R_g$ | $R_g^2=\frac{1}{N}\sum_{i=1}^{N}\lvert\mathbf{r}_i-\mathbf{r}_{\text{cm}}\rvert^2$ |
| End-to-end distance | $R_{ee}$ | $R_{ee}=\lvert\mathbf{r}_N-\mathbf{r}_1\rvert$ |
| Hydrodynamic radius | $R_h$ | Kirkwood approximation $\left\langle\frac{1}{R_h}\right\rangle=\frac{1}{N^2}\sum_{i\neq j}\frac{1}{r_{ij}}$ |
| Normalized number of contacts | $N_{\text{contact}}^{\text{norm}}$ | $\frac{N_{\text{contact}}}{N}$ |

Table S4: Proteins used for ortholog molecular dynamics (MD) simulations

| **Cluster ID** | **UniProt ID** | **Residue Range** | **Number of Orthologs** |
|---|---|---|---|
| 1 | O14686 | 3966–5024 | 123 |
| 1 | P29400 | 1–1456 | 107 |
| 1 | Q01955 | 1–1441 | 114 |
| 1 | P08123 | 1–1134 | 193 |
| 1 | Q9UPS6 | 897–1823 | 178 |
| 1 | Q14031 | 1–1464 | 113 |
| 1 | P05997 | 85–1266 | 198 |
| 1 | P02458 | 78–1254 | 220 |
| 1 | P20908 | 242–1612 | 192 |
| 1 | Q9UMN6 | 1781–2416 | 106 |
| 1 | Q9UMN6 | 1–966 | 97 |
| 1 | P13942 | 219–1546 | 178 |
| 1 | Q8NEZ4 | 460–946 | 152 |
| 1 | P08572 | 1–1484 | 152 |
| 1 | Q8NEZ4 | 1782–3172 | 158 |
| 1 | P02462 | 1–1436 | 179 |
| 1 | P02461 | 79–1230 | 98 |
| 1 | P12107 | 243–1579 | 197 |
| 1 | Q8NEZ4 | 3423–4385 | 158 |
| 1 | O14686 | 2704–3505 | 110 |
| 1 | O14686 | 345–1364 | 121 |
| 1 | O15047 | 829–1417 | 58 |
| 1 | Q8NEZ4 | 1–234 | 133 |
| 2 | P22105 | 2070–2101 | 34 |
| 2 | P22105 | 2275–2308 | 31 |
| 2 | Q6NUI6 | 720–762 | 115 |
| 2 | P20336 | 190–220 | 241 |
| 2 | P22105 | 1345–1379 | 45 |
| 2 | P22105 | 2710–2742 | 22 |
| 2 | Q9NRW1 | 176–208 | 164 |
| 2 | P22105 | 3035–3066 | 2 |
| 2 | P22105 | 1962–1992 | 29 |
| 2 | P22105 | 2495–2528 | 28 |
| 2 | P22105 | 2386–2415 | 18 |

*Continued on next page*

Table S4 – *Continued from previous page*

| Cluster ID | UniProt ID | Residue Range | Number of Orthologs |
|---|---|---|---|
| 2 | P20340 | 177–208 | 178 |
| 2 | P22105 | 1857–1888 | 36 |
| 2 | P51159 | 188–221 | 199 |
| 3 | Q96HR8 | 1–171 | 169 |
| 3 | Q8IU60 | 242–420 | 177 |
| 3 | O14746 | 174–331 | 171 |
| 3 | P04259 | 1–161 | 112 |
| 3 | P13647 | 1–166 | 95 |
| 3 | P12035 | 508–628 | 60 |
